# Crowdsourcing Machine Intelligence Solutions to Accelerate Biomedical Science: Lessons learned from a machine intelligence ideation contest to improve the prediction of 3D domain swapping

**DOI:** 10.1101/2020.07.12.199398

**Authors:** Yash Shah, Deepak Sharma, Rakesh Sharma, Sourav Singh, Hrishikesh Thakur, William John, Shamsudheen Marakkar, Prashanth Suravajhala, Vijayaraghava Seshadri Sundararajan, Jayaraman Valadi, Khader Shameer, Ramanathan Sowdhamini

## Abstract

Machine intelligence methods, including natural language processing, computer vision, machine vision, artificial intelligence, and deep learning approaches, are rapidly evolving and play an essential role in biomedicine. Machine intelligence methods could help to accelerate image analyses aid in building complex models capable of interpretation beyond cognitive limitations and statistical assumptions in biomedicine. However, irrespective of the democratization via accessible computing and software modules, machine intelligence handiness is scarce in the setting of a traditional biomedical research laboratory. In such a context, collaborations with bioinformatics and computational biologists may help. Further, the biomedical diaspora could also seek help from the expert communities using a crowdsourcing website that hosts machine intelligence competitions. Machine intelligence competitions offer a vast pool of seasoned data scientists and machine intelligence experts to develop solutions through competition portals. An alternate approach to improve the adoption of machine intelligence in biomedicine is to offer machine intelligence competitions as part of scientific meetings. In this paper, we discuss a structured methodology employed to develop the machine intelligence competition as part of an international bioinformatics conference. The competition leads to developing a novel method through crowdsourcing to solve a challenging problem in biomedicine – predicting probabilities of proteins that undergo 3D domain swapping. As a biomedical science conference focused on computational methods, the competition received multiple entries that ultimately helped improve the predictive modeling of 3D domain swapping using sequence information.

## Background

The rapid advent of advanced molecular profiling and experimental methods, including sequencing, gene-editing, and multi-omics technologies during the last decade, has enabled biology to enter the era of ‘Big Data.’(1–8). However, the computational efficiency of analyzing and interpreting petabyte-scale data has become a bottleneck. Although significant “interpretation gap” in biomedicine where extensive, multi-scale data on various disease modalities exist, the collective impact of defining such datasets remain limited. In this regard, bioinformatics approaches have uprooted wherein robust statistical learning and reproducible machine intelligence methods are evolving to play a crucial role in addressing the inference challenges.

In this era of smart algorithms and artificial-intelligence driven knowledge banks, bioinformatics applications could make an immediate impact in the setting of molecular medicine, drug development, crop improvement, gene therapy, microbial genome annotation, and assembly, etc. Integrating the complexity of biomedicine data with modern machine intelligence methods followed by orthogonal and experimental validations could lead to uncovering new biological themes and ultimately aid in discovery. The recent advances in different areas of machine intelligence, including deep learning, reinforcement learning, and growth towards developing general intelligence, would eventually enable such methods as a pivotal part of biomedical discovery research(7,9–11).

## Challenges in democratizing machine intelligence for biomedicine

Machine intelligence methods are currently going through an “AI Spring” with extensive focus on developing new techniques. Industry sectors across different verticals, including healthcare, life science, biotech, pharma, and medical device technology is making significant investments to improve data access along with design, development, and deployment of machine intelligence methods(12–14). Design development and implementation of reproducible machine intelligence approaches need substantial investment to acquire diverse talent, computing resources, and deployment infrastructure.

To illustrate the complexity of one factor: computing resourcing– extensive evaluations are often required to leverage in-house, cloud, or hybrid mode of computing infrastructure(15). Depending on the nature of the problem to solve, computing infrastructure and software systems could be selected. For example, cloud computing vendors like Microsoft Azure, Google, Amazon Web Services offers a wide variety of operating systems, database solutions, operating systems, and machine learning frameworks along with a custom framework for rapid design, development, and deployment of machine learning solutions. Ultimately, the choices and decisions in every step could influence the cost of computing and the development of machine intelligence solutions. Delivery of a machine intelligence solution requires a team that comprises a domain expert to curate and interpret the data, data engineer to clean and compile data sets and data scientists to develop the model. Often the implementation needs solution architectures and software engineers to build web service, client-server architecture, and endpoints as web or mobile applications(16).

### Crowdsourcing, Online Competitions and Innovation Contests: Past, Present and Emerging Trends

Online coding competitions, such as Kaggle data competitions, Netflix data competitions, Google Code Jam, all help participants practice critical thinking, fast and efficient coding, and the ability to design and then implement algorithms in code. These competitions allow for coders to learn new technologies that they would not have learned otherwise. These new technologies and techniques can then be brought back to the coder’s daily lives at work for improvements and to overcome obstacles in a unique, refreshing manner. These competitions are grounds for new and different approaches to solving a common problem, by experiencing more ways to tackle a problem, participants can learn coding standards and adapt methods proposed by creative coders, with more tools at hand. Furthermore, the issues at various online coding competitions are modeled after real-world problems. For example, Kaggle, a highly popular site that hosts data science and machine learning competitions, provides data sets that expose participants to forecasting, sentiment analysis, natural language processing, and image classification problems. These topics are very applicable to real-life problems and are also at the forefront of current technology. Although there is a wide range of topics, each question that a participant picks forces them to learn about the context of the problem, the data, and the approach to the problem. Of course, this will spawn many nuanced different techniques that everyone can view and glean information from.

Competitions at their heart compare the outputs of different teams, and a ranking system spurs on coders to become better, more efficient, and they are an excellent way for a bioinformatician, data scientist, data engineer or software developer collectively called as a coder to test themselves based on the population. With official online competitions, participants have to work efficiently and on-time, while making sure their solutions are time and memory efficient. With competition, there is a constant need for the participant to improve their code and look for ways to separate themselves from the pack. If everyone has the same data and access to algorithms, coders are forced to find ways to push their code forward, which will lead to innovations. While people find success in these online competitions, other coders that are looking to begin participating in coding competitions can look up to top performers as role models, and experts in the respective fields will guide them by following the winning code. The desire to win a competition, will create changes in the field of study, push the previously established limits of performance, and also connect like-minded or different-minded people to form communities that will be able to tackle the problem in many different ways.

For example, the ImageNet challenge is an example that revolutionized the domain of deep learning applied to computer vision(17). ImageNet challenge, which revolutionized the field of deep learning and applied computer vision. ImageNet Large Scale Visual Recognition Challenge (ILSVRC) started in 2010 with a dedicated database for visual recognition called ImageNet dataset, which is also a result of crowdsourcing. The image-level annotation of the database is done by crowdsourcing with more than 14 million annotated images for visual/ object recognition. It was in 2012, Alex Krizhevsky came up with a model named AlexNet(18), which performed incredibly in the contest with a top-5 error of 15.3% and the manuscript “Imagenet classification with deep convolutional neural networks” has got more than forty thousand citations till date. AlexNet introduced many new methods, including the GPU utilized training, which fueled the deep learning and computer vision revolution. After AlexNet, it was a rally of algorithms and architectures with better performance in the followed years’ contests, including Microsoft’s ResNet(19) and inception by Google. Several other machine learning and deep learning contests and challenges emerged from the inspiration of the Imagenet problem in recent years. And all of them, students, and researchers across the globe are contributing to the community in association with these challenges.

## Crowdsourcing machine learning solutions in biomedicine

Crowdsourcing is the practice of engaging a ‘crowd’ or group for a common goal to innovate, design, solving a problem(1,20). With a lot of unsolved problems in biology, the use of crowdsourcing to solve important but complex problems in biomedical and clinical sciences is growing and encompasses a wide variety of approaches which include data mining crowd-generated data in healthcare or open source platforms [4]. To democratize machine intelligence and familiarize the research community with machine intelligence methods, crowdsourcing competitions could be an ideal solution. Crowdsourcing is emerging as a recent trend in biomedical science that aims to tap into the skills not immediately available in a laboratory setting due to specialtyor scalability of a task.One of the classical examples of crowdsourcing in biomedicine includes Folding @Home, which aim to use idle computing time from registered users to perform computationally expensive protein folding classification. Further, biomedical applications that benefitted from crowdsourcing includes genomic variant curation, bioinformatics research, health surveillance, protein folding research, proteomics, environmental research, stem cell biology research, public health research and data visualization (See **Table: 1**)(21–42). Recent examples including classification of acoustic datasets, identification of chemical induced diseases, clinical trial result summarization, therapeutic area-specific knowledge assimilation in the area of dermatology and plant phenomics (43–47). A conceptual framework for crowdsourcing an ideation contest is given in **Figure: 1**.

**Figure 1:**
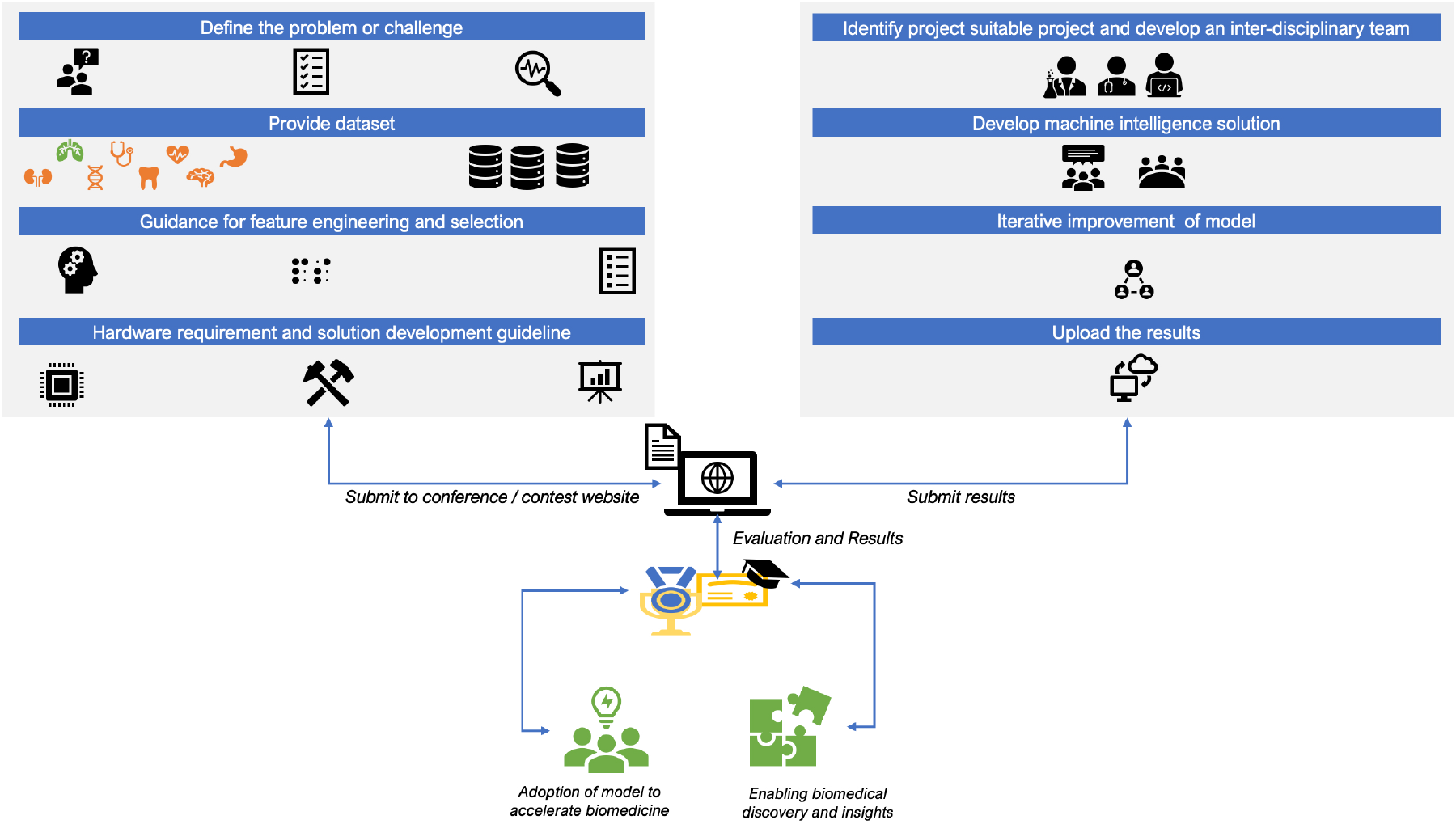
Strategic framework for developing and organizing an ideation contest

**Table 1:**
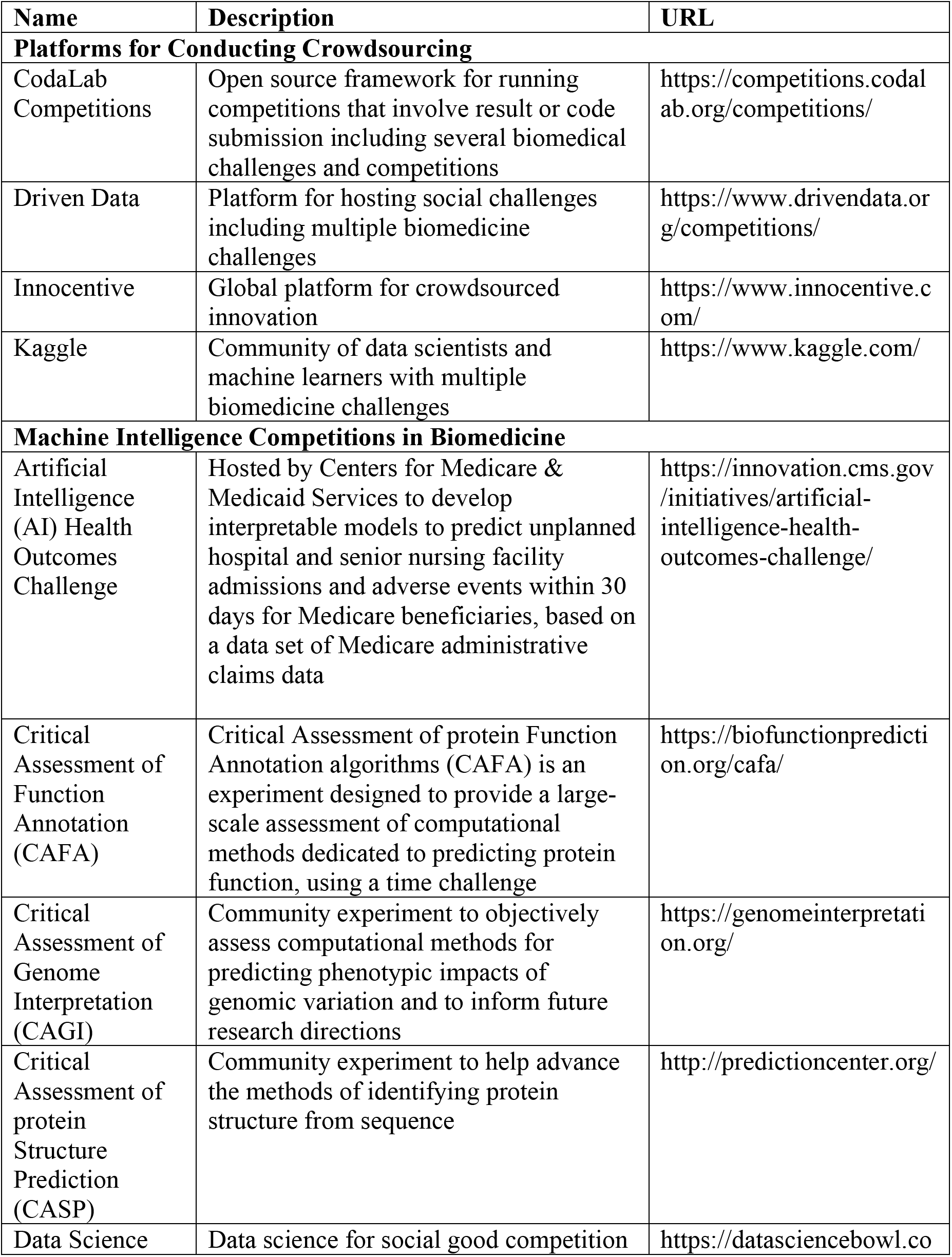
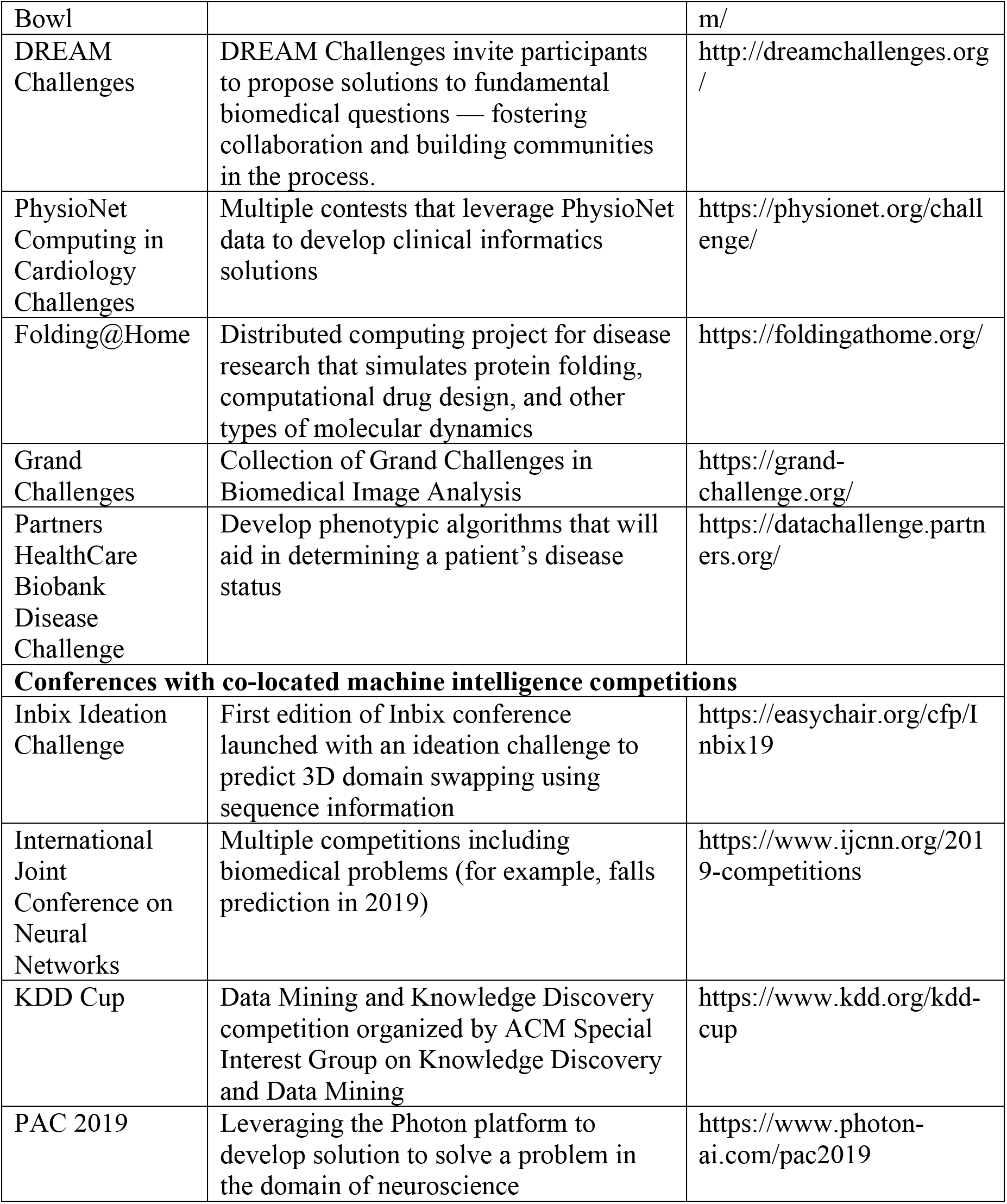
Competitions, Ideations, Conferences and Platforms for crowdsourcing in biomedicine

## 3D Domain Swapping

3D domain swapping is a mechanism by which two or more protein chains form a dimer or higher oligomer by exchanging an identical structural element. While the mechanism was first observed in 1964 and conceptualized in 1994(48–51). Proteins, including antibody fragments, human prion protein, crystallins, growth factors, cytokines, etc. are involved in 3D domain swapping. The precise roles of domain swapping as a causal factor of different disease pathways, including conformational and deposition diseases, to remain elusive(52). However, experimental studies have suggested that change in environment (low pH, temperature, denaturants) or genetic predisposition may lead to 3D domain swapping. A systematic survey of 293 proteins with swapped conformation revealed several biological clues including the functional landscape, disease associations and pathways that are driven by proteins in swapped conformation(53). Biophysical impact, including the kinetic effect (closed interface) or dynamic effect (open interface), has also been suggested. A curated knowledgebase of proteins involved in 3D domain swapping “3DSwap” is available in the public domainfrom http://caps.ncbs.res.in/3dswap/based on the graduate research by one of the Ideation contest developer (KS) and supervised by the Ideation contest evaluator (RS)(54). Data compiled in 3DSwap database was used to establish first prediction algorithms using machine learning and artificial intelligence approaches including support vector machines (SVM; model accuracy 63.8%) and RandomForest (RF; model accuracy: 73.81%) models(55,56). These models can perform prediction, instead of experimental characterization of domain swapping. Where the latter is expensive and time-consuming, prediction algorithms were applied to human proteome and identified new proteins to be associated with features of swapping.

### Crowdsourcing to Improve Prediction of 3D Domain Swapping from Sequence Information

Indian Conference on Bioinformatics (Inbix’17) held at Birla Institute of Scientific Research. Jaipur, India. The Inbix’17 program had a participation of 190 delegates besides keynote speakers, invited speakers, oral and poster, and ideation challenge presenters. We asked the Inbix’17 conference attendees to improve this sequence-based model published in 2010/2011 and provide a higher accuracy model by adopting new feature engineering strategies and novel machine learning approaches, including deep learning. The model with better accuracy and biologically relevant feature engineering approached was highly encouraged as part of the results submission.

### Guidelines for contest to improve prediction of 3D domain swapping

Problem definition (See Supplemental Material for Ideation Challenge notice) and link to access data set (See Supplemental Material for positive and negative datasets) was given to the participants of the conferences using the website of the meeting. Conference organizers used social media and other outlets to publicize the contest across the world. No additional guidelines were given to generate features or the selection of machine learning, as this may hinder novel contributions from the community. The results were compiled using an evaluation framework by a team of researchers. Models were evaluated for innovation, feature engineering strategy, algorithm applied to develop predictive model, the robustness of validation method, and net improvement in the prediction of 3D domain swapping compared to the model published earlier.

### Proposed solutions

Each of the three predictive models proposed approached the problem in different ways (Also See **Table: 2**). Preprocessing techniques like feature extraction and feature selection are way different, and a variety of model optimization methods are tried to get a more accurate prediction model possible. A brief description and critical appraisal of the models are given below for brevity. A fourth submission was a conceptual overview to address the biological knowledge gap in the setting of 3D domain swapping. The original version of the submission of the solution for different solutions and all associated data and code is available in the Supplemental Materials (Also see **Figure: 2**).

**Figure 2:**
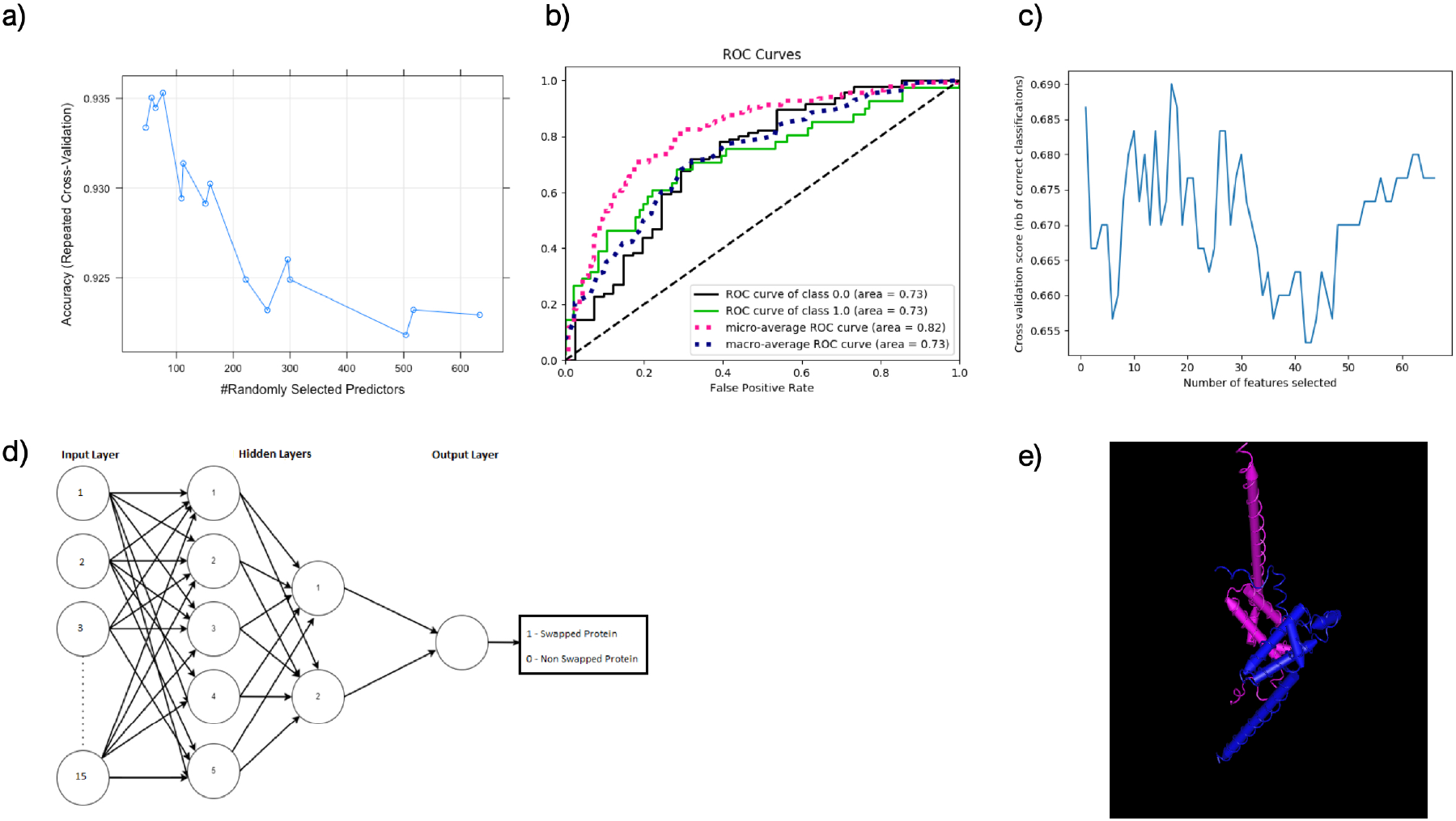
Summary of machine intelligence strategies used to improve prediction of 3D-domain swap using machine intelligence methods. a) Plot between Accuracy and number of randomly selected predictors used for grid searching in Model-1; b) ROC curve of Model-2; c) Features selected v/s cross-validation scores on training samples compiled from Model-3. d) Neural network architecture used in Model-2 e) 3D model of Ubiquitin carboxyl-terminal hydrolase 8 – a human hydrolase enzyme.

**Table 2:** Summary of models submitted to ideation contest to improve the prediction of 3D domain swapping from sequence information

| **Models** | **FE-Strategy** | **Algorithm** | **Reported AUC** | **Packages** | **Features** |
| --- | --- | --- | --- | --- | --- |
| Model-1 | Boruta method | Nnet | 90.73% | Boruta, nnet,<br>neuralnet | 8521 |
| Model-2 | Mutual<br>information gain | XGBoost | 75.63% | Scikit-learn,<br>XGBoost | 369 |
| Model-3 | Selectkbest | MLP | 72.5% | scikit-learn | 66 |

### Summary and Critical evaluation of Models

#### Model 1

The first model coded entirely in R language, uses R library packages for both feature engineering and model design. The preprocessing is a crucial part of machine learning, which includes data cleaning, feature extraction, and feature selection. R package named “peptides” (See: https://cran.r-project.org/web/packages/Peptides/index.html) is doing all the feature generation jobs in this model. Peptides package has several useful functions to calculate indices and physicochemical properties of protein sequences. Boruta package in R was used for feature selection (57), which is the process of selecting only the relevant features which affect domain swapping, and 652 features are finally confirmed. Boruta uses Random Forest by default to search for relevant features by comparing primary attribute importance with importance achievable in random, subtracting the irrelevant features to stabilize the needed features. Even though time-consuming, the boruta method outputs enough, and efficient data for the model to learn, gives the best model accuracy and avoids the problem of overfitting. The final dataset with total samples of 1185 is split cleverly into 80%: 20% for training and testing, respectively. This data is trained on a random forest algorithm with a set of well-tuned parameters. Successful optimization for parameters like the number of tress and mtry gives a final result of 91.03% training accuracy, 91.22% accuracy after five-fold cross-validation, and 90.29% testing accuracy with 3000 trees and 656 mtry. The R package *Boruta* was used for feature extraction. The modelers split that into the ratio of 80:20 for training and test set. No validation set was provided. The submitters used the training set to select models, which are slightly flawed as there may be a possibility of the model overfitting on the training set and them not knowing until they do their final test on the test set. Note that a shallow neural net and various R neural net packages did not perform as well as the random forest model used by the earlier model. This may be due to the nature of the data, data preprocessing, not enough resources or time to train a massive neural net, or just a flawed implementation of the correct network structure. The group achieved a somewhat high training and validation accuracy of around 90%, much higher than the other groups.

#### Model-2

The second model proposes a method called ensemble modeling in which soft voting is carried out between two classifiers after feature engineering. In the feature engineering part, efficient data cleaning is done as the redundant sequences are entirely dropped. Concentrating more on data cleaning and feature engineering, this model uses different library packages for feature extraction like modlAMP (See: https://modlamp.org/), etc. Feature selection is carried out by using python’s scikit-learn’s simple but powerful feature selection library called selectKbest, which uses mutual information gain for selecting top features(58,59). An ensemble model is proposed in which soft voting is carried out between two simple and accurate predicting algorithms, AdaBoost and XGBoost(60). Both the algorithms are fine-tuned to get the best parameter possible with the data. The fine-tuned models are put in a Voting Classifier with a weight of 4:6 with a majority in favor of XGBoost Classifier. A better performance was observed with an accuracy of 75.63% after five-fold cross-validation.

#### Model-3

While the first two models focus on solving the problem using classical machine learning algorithms, the third model uses an artificial neural network algorithm, which comes under the deep learning approaches. Standardization is done on the features to make all the features on a common scale with zero mean and unit variance. This ensures less computation time and removal of data overfitting by bringing the range and scale of the feature variables to a standard measure. Especially in multi-layer perceptron (MLP(61)) models, standardization is usually done on the data to decrease the time taken by the model for weight optimization. The feature selection part removes redundant features and reduces the dimensionality of the dataset to ensure reasonable accuracy and an improved result. Feature selection model selected top 15 features out of 66 features extracted from the training and testing phase of the modeling. The conventional train test split of 70:30 is done on the sample of size 426, which gives 300 examples for training and 126 samples for testing. The train-test data is well-balanced such that both of them consist of half of the samples from every two classes. That is, 300 training samples have 150 samples from positive class and 150 from negative class in it, to avoid class imbalance. The data is then fed into an MLP, which is an artificial neural network classifier that uses back propagation algorithm for learning and error correction. The model follows simple multi-layer neural network architecture with five neurons in the first hidden layer and two neurons in the second hidden layer. Hyperparameters like a number of hidden layers, activation function, and solver are optimized and fine-tuned to reach out to the best result of 76.67% accuracy in 10-fold cross-validation and test accuracy of 72.5%. The coding part is supported by several python-machine learning library modules from scikit-learn such as SelectKbest for feature selection, MLP, StandardScaler for data normalization, and other modules for metrics and cross-validation. These packages help to implement a useful model in a few lines of code. One problem may be the lack of training examples; only 150 positive and 150 negative data points were used. The network was not very deep as it was two layers deep with five and then two units, respectively considering limited training data set. This was an interesting approach as it attempted to use a neural network to approximate a nonlinear function. However, many questions arise from this implementation: including the need for a deep learning approach is necessary or overengineering the problem. It is unclear whether the proposed layers are enough or more hidden layers, and units are needed to learn a machine problem with a limited dataset. Alternate network structures that could work better than a feed-forward network was not addressed.

## From domain swapping to drug targeting: pushing the boundaries on targeting domains of unknown function

The 3D swap database has a couple of domains of unknown function (DUF), which we would like to consider a case study to infer the role of aptamers. Assuming that the functions of DUFs and hypothetical proteins (HP) can leverage as targets for diagnostics, the most common entity used are antibodies which could circumvent the effect/targets. While the experimental characterization of antibodies is cumbersome, it is assumed that aptamer-protein prediction methods may serve as a benchmark besides providing cost-effective measures(62–64). In this ideation example, we propose a hypothesis whether the aptamer is bound in the setting of a 3D domain-swapped conformation. If so, could it be applied for domains caused due to extensive multimerization as well? To answer this, we have considered the DUFs with a PDB entry 2A9U (http://caps.ncbs.res.in/cgi-bin/mini/databases/3Dswap/3dswap_entry.cgi?id=2A9U and **Figure 2**). As there is a dimer interface communicated to the catalytic domain of 2A9U, we assumed that the aptamers specific to this variable fragment could be used. With this approach, we expect that through the antigen-binding capacity of aptamer with the molecule, a vast number of HPs or DUFs can be targeted, which could be associated with diseases. Thus, authors hope active conformation and aptamers as small molecules for therapies could prove to be very useful in the development of treatment for several diseases where 3D domain swapping is a known pathological mechanism. To conclude, we hypothesize that the role of aptamers over antibody isotypes can be inferred and based on the affinity of aptamers bound to swapped domains particular to HPs or DUFs.

## Discussion

With the current status of poor outcomes in recent clinical trials in the setting of neurodegenerative diseases, novel drug discovery and drug repositioning approaches are required to address the pathological basis of protein conformation diseases like Alzheimer’s diseases(65–69). Collectively, the ideation contest helped to apply modern algorithms, new feature engineering and feature selection methods to enhance the prediction of 3D-domain swapping – a key mechanism in the setting of conformational diseases. Improving the prediction accuracy of 3D domain swapping from sequence information using machine learning is critical to enable the rapid characterization of a novel structural phenomenon. In this paper, we discuss about developing an ideation contest to improve prediction of 3D domain swapping from sequence information. We discuss about the 3D domain swapping mechanism and provide an overview of model proposed by leveraging different machine learning approaches to predict whether a given protein is swapping or non-swapping. 3D domain swapping is a process through which a protein oligomer is formed from their monomers.

The rationale for predict 3D domain swapping from sequence information is based on the classical Anfinsen’s dogma postulation that the native structure of a protein sequence is determined by the properties of the amino acids of that protein sequence. Three different machine learning approaches were proposed by the ideation contestants for successfully predict and classify proteins into swapping or non-swapping proteins.Compared to the original models published in 2010 and 2011; modern machine intelligence approaches helped to improve the model modestly. The improvement could have been much better with more data availability. Thus, proposing machine intelligence contests as part of biomedical conferences may help to enhance the discovery of novel biomedical insights.

## Conclusions

Biomedical Data Scientists could design and develop Machine Intelligence-based informatics solutions to address challenges in biology and medicine. Machine Intelligence is evolving as a critical analytical theme in biomedicine due to the advent of big data, scalable and affordable cloud computing and modern machine learning toolkits. Leading biomedical science and informatics conferences could organize ideation contests, predictive modeling competitions and crowdsourcing efforts to improve the democratization of machine learning in bioscience. We used a machine learning ideation competition to revisit the problem of predicting the 3D domain swapping - a mechanistic basis of protein conformations in neurodegenerative diseases; as part of an international bioinformatic conference. New insights and a variety of solutions were proposed to address the challenging problem of predicting protein aggregation mechanism from sequence information. Collectively, the crowdsourcing results from ideation competition could help to push the conceptual boundaries and unlock new ideas to understand complex mechanisms like 3D domain swapping.

## Data, Source Code and Model Availability

- Supplemental Materials are available from figshare: https://doi.org/10.6084/m9.figshare.8317067.v1

Source code is available at the following repositories:

○ Model-1: https://github.com/DBT-BIF/Inbix_ideation
○ Model-2: https://github.com/souravsingh/Ideation-Challenge
○ Model-3https://github.com/shahyash-95/ideation.challenge_inbix2017

## Acknowledgements

R.S. and K.S. acknowledge National Centre for Biological Sciences (TIFR) for infrastructural and financial support. R.S. was a Senior Research Fellow of the Wellcome Trust, U.K. R.S. and G.A. thank Department of Biotechnology, Government of India for financial support. Rakesh and Deepak acknowledge the infrastructure support from the Bioinformatics Infrastructure Facility (DBT-BIF), University of Rajasthan, Jaipur.

## Biographical Notes

Yash Shah holds a Bachelor’s degree in Computer Engineering from Mumbai University, an experienced software engineer and currently working as Research Bioinformatician at ACTREC, Tata Memorial Centre, Mumbai, Maharashtra, India.

Deepak Sharma is a Doctoral student in Indraprastha Institute of Information Technology, Delhi, and a Senior Research Fellow in the Institute of Nuclear Medicine and Allied Sciences (DRDO), New Delhi, India.

Rakesh Sharma is a Bioinformatician in Bioinformatics Infrastructure Facility, University of Rajasthan, Jaipur, India.

Sourav Singh has a BE degree in Computer Engineering from VIIT, Pune, India.

Hrishikesh Thakur holds an M.Tech Degree in Modelling and Simulation from Savitri Bai Phule Pune University, Pune, India.

William John is an alumnus of the Computer Science Department, New York University, New York, NY, USA.

Shamsudheen Marakkar is a student a Robotics and Artificial Intelligence MTech student at Cochin University of Science and Technology, Kochi, Kerala, India.

Prashanth Suravajhala is a Senior Scientist in Systems Genomics based in Birla Institute of Scientific Research, Jaipur, India. He can be reached at http://wiki.bioinformatics.org/prash

Vijayaraghava Seshadri Sundararajan is a Data Scientist in Singapore.

Jayaraman Valadi is currently a Distinguished Professor of Computer Science at Flame University, Pune, India.

Khader Shameer was a member of the Institute for Next Generation Healthcare, Icahn School of Medicine at Mount Sinai, Mount Sinai Health System. At the time of publication, Shameer is a Senior Director (Data Science, Advanced Analytics, and Bioinformatics) with AstraZeneca.

Ramanathan Sowdhamini is a professor at the department of biochemistry, biophysics, and bioinformatics of the National Centre for Biological Sciences and leads the Computational Approaches to Protein Sciences laboratory.

