## Supplementary material for "Crowdsourcing Machine Intelligence Solutions to Accelerate Biomedical Science: Lessons learned from a machine intelligence ideation contest to improve the prediction of 3D domain swapping": Inbix_Ideation_Evaluation_Form_Template.docx

Inbix Ideation contest details: <https://drive.google.com/file/d/0Bxue5qHTjmJIc2hieVZZbVBnNnM/view>

Each entry will be assessed for **5 parts**; each section carrying **10 points each**.

Ideally, the best model will receive a score of 50 points.

a. Innovation – overall innovation adapted by the team

b. Feature engineering strategy

c. Strategy for selection of prediction algorithm

d. Validation approaches

e. Improving current prediction accuracy (Sequence-based: 62.34; Sequence and Structure-based prediction: 73.81)

| Name | Innovation | FE-Strategy | Algorithm | Validation | Improvement |
| --- | --- | --- | --- | --- | --- |
| Model 1 |  |  |  |  |  |
| Model 2 |  |  |  |  |  |
| Model 3 |  |  |  |  |  |
