## Supplementary material for "Crowdsourcing Machine Intelligence Solutions to Accelerate Biomedical Science: Lessons learned from a machine intelligence ideation contest to improve the prediction of 3D domain swapping": Inbix_Invitation_Ideation_Challenge_ML_v1.pdf

Indian Conference

on Bioinformatics 2017

### Improving the prediction accuracy of 3D domain swapping from sequence information using machine learning

3D domain swapping is a mechanism by which two or more protein chains form a dimer or higher oligomer by exchanging an identical structural element. While the mechanism is observed in 1964 and conceptualized in 1994, its significant role in disease pathophysiology was unclear. Proteins including antibody fragments, human prion protein, crystallins, growth factors, cytokines, etc. are involved in 3D domain swapping. The precise roles of domain swapping in causing different disease pathways to remain elusive. However, some of the experimental studies have suggested that change in environment (low pH, temperature, denaturants) or genetic predisposition may lead to 3D domain swapping. Biophysical impact including kinetic effect (closed interface) or dynamic effect (open interface) has also been suggested. A curated knowledgebase of proteins involved in 3D domain swapping “3DSwap” is available in the public domain.  Data compiled in 3DSwap database was used to establish first prediction algorithms using machine learning and artificial intelligence approaches including support vector machines (SVM; model accuracy 63.8%) and RandomForest (RF; model accuracy: 73.81%) models. These models can now perform prediction, instead of experimental characterization of domain swapping. Where the latter is expensive, laborious and time-consuming. Prediction algorithms were applied to human proteome and identified new proteins to be associated with features of swapping. We invite the Inbix ’17 conference attendees  to improve this sequence-based model published earlier and provide a higher accuracy model by adapting modern feature engineering strategies and novel machine learning approaches including deep learning. The model with better accuracy and biologically relevant feature design plans are highly encouraged.

**Rules of the ideation challenge:**

1. A team per institution, minimum 2 members and maximum 5 members per team including students, interns, employees or faculty members
2. Authors should use the data from 3Dswap database, develop new sequence and/or structure feature engineering strategies and develop a new model
3. Authors should report training, testing and 5-fold cross-validation accuracies
4. Annotated code related to the feature engineering and machine learning should be provided in Python or R. If a GUI based package (Weka, Orange etc.) is used for deriving features or creating models – a copy of the same should be provided to for evaluation
5. Each entry will be assessed for 5 parts; each section carrying 10 points each. Best model will receive a score of 50 points.
    a. Innovation – overall innovation adapted by the team
    b. Feature engineering strategy

c. Strategy for selection of prediction algorithm
d. Validation approaches
e. Improving prediction accuracy

**Data set:**

Positive data set: http://caps.ncbs.res.in/3dswap-pred/data/3dswap-pred_positive_dataset.fasta
Negative data set: http://caps.ncbs.res.in/3dswap-pred/data/3dswap-pred_negative_dataset.fasta

**Format for reporting results – Ideation document:**

- Name of the team, affiliations, email and contact numbers
- Data summary (positive data, negative data). Teams may improve the positive or negative data; all data wrangling steps should be explained in the ideation document
- Feature engineering: provide detailed account of feature engineering, feature selection strategies
- Model summary: motivation behind selection of model, model accuracies and validation
- Biological summary: relevance of the model in the biological or clinical context.
- Link to github/bitbucket with code

**Related Readings.**


3DSwap: curated knowledgebase of proteins involved in 3D domain swapping
K Shameer, PN Shingate, SCP Manjunath, M Karthika, G Pugalenthi, R Sowdhamini
Database 2011

Insights into protein sequence and structure-derived features mediating 3D domain swapping mechanism using support vector machine based approach
K Shameer, G Pugalenthi, KK Kandaswamy, PN Suganthan, G Archunan, R Sowdhamini
Bioinformatics and biology insights 4, 33

3dswap-pred: Prediction of 3D Domain Swapping from Protein Sequence Using Random Forest Approach
K Shameer, G Pugalenthi, K Kumar Kandaswamy, R Sowdhamini
Protein and Peptide Letters 18 (10), 1010-1020

Functional repertoire, molecular pathways and diseases associated with 3D domain swapping in the human proteome
K Shameer, R Sowdhamini
Journal of clinical bioinformatics 2 (1), 8

**Ideation challenge coordinators:**
Dr. Khader Shameer and Dr. V.S Sundararajan

**Ideation challenge evaluators:**
Dr. Ramanathan Sowdhamini, Dr. Atul K Upadhyay, Dr. Khader Shameer and Dr. V.S Sundararajan

**How to Submit?**
Please submit through easychair.
