## Supplementary material for "Crowdsourcing Machine Intelligence Solutions to Accelerate Biomedical Science: Lessons learned from a machine intelligence ideation contest to improve the prediction of 3D domain swapping": Model-1_RakeshSharma_Inbix17_paper_52.pdf

### Inbix-2017 ideation challenge report


#### Team name: DBT-BIF

Name: **Rakesh Sharma**

- Affiliations: BIF (Bioinformatics Infrastructure Facility), University of Rajasthan
-
- Contact no.: +91-8955464254

Name: **Deepak Sharma**

- Affiliations: Institute of Nuclear Medicine and Allied Sciences
-
- Contact no.: +91-9602431619


#### Data summary

We used R for feature engineering, feature selection and data modeling. Positive and negative dataset were merged into one dataset for the sake of easy handling of data. This initial dataset contains peptide sequences that contain amino acids other than 20 naturally present amino acids. Our solution comprised of data wrangling, we removed peptides sequences that contain non natural amino acids. All the non-conforming records (other than 20 default types) were rejected. We also rejected peptide sequences having length less than 10 amino acids, since such peptides sequences are non-functional in nature. There were no missing values, so no 'missing value treatment' was given. Final dataset contains 728 positive swapping cases and 457 negative cases.

#### Feature engineering

Feature generation:

R package 'peptides' was meticulously used to generate biologically significant features. Initially basic features such as properties of amino acid (Tiny, Small, Aliphatic, Aromatic, Nonpolar, Polar, Charged, Basic, Acidic, etc.) were used. We pondered on other biological properties that affect domain swapping. Amino acids dimers and trimers were supposed to add swapping functionalities in peptide sequences. Thus we generated dimers and trimer permutations of all naturally occurring amino acids.

Feature selection:

Of several feature selection strategies, ‘Boruta’ feature selection method provided the best results. According to R ‘Boruta’ package – “Boruta finds relevant features by comparing original attributes' importance with importance achievable at random, estimated using their permuted copies”. Boruta wrapper algorithm results in categorization of all features in 3 classes – ‘confirmed important’, ‘tentative’ and ‘confirmed unimportant’. We selected only those features that are confirmed to be important.

#### Model summary

We used supervised learning approach for this classification problem.

As mentioned above our final dataset contains 1185 total cases, after splitting dataset of these 1185 cases, 948 (80%) cases were used for training the various models and 237 (20%) cases were used to test the model accuracy.

Search for best classification model started with training our data with multilayer perceptron model using R’s **nnet** package (1 hidden layer) and **neuralnet** package with various neural network architecture but the test case accuracy wasn’t quite satisfactory. This search moved on to ‘Random Forest’ model and as expected ‘Boruta’ selected features performed well on test case and their cross-validation accuracy was impressive as well. Random forest model for classification was implemented in R using RandomForest package.

Our model was trained and tested using 3000 trees, 656 mtry having maximum possible 450 nodes. Other important parameters are class weight [0.385,0.614] and we took 300 samples from each class, out of bag times parameter was set to 1000 with node size 2.

Training accuracy for random forest model was 90.40%, validation accuracy was 90.50% and 5-cross validation accuracy calculated was 90.73%.

#### Biological summary

We figured out that basic properties of amino acids such as physical and chemical properties impact directly on domain swapping and that dimer, trimer sequence hardly have any significance in domain swapping. We observed that using these simple properties of amino acids one can develop a classification model.

#### Feature  Link to source code (Github)

This work was accomplished using Github as a version control system using R language, for which source repository is available on following link.

https://github.com/DBT-BIF/Inbix_ideation
