## Supplementary material for "Crowdsourcing Machine Intelligence Solutions to Accelerate Biomedical Science: Lessons learned from a machine intelligence ideation contest to improve the prediction of 3D domain swapping": Model-2_SouravSingh_Inbix17_paper_60.pdf

### Solution for Ideation-Challenge

#### The document summarizes the approach used for solving the problem of obtaining a higher prediction accuracy for 3-D domain swapping using Machine Learning methods.

#### Data Cleaning and Feature Engineering method

The data for the challenge was extracted from here- Positive dataset and Negative dataset. Upon extraction of the dataset, Redundant Sequences from the positive FASTA dataset was removed using CD-HIT from Fu et al. 2012 at 95% cut-off ratio. For Negative FASTA sequence dataset, no redundancy removal was done, but sequences containing non-natural amino acids like the letter 'X' were removed from the dataset, which constituted 5 out of 462 negative FASTA sequences.

Once the data cleaning was complete, Feature extraction for the Positive and Negative FASTA sequences was done using the modlAMP package from Müller et al. 2017, from which non-amino acid scale dependent features like Molecular Weight, Charge Density, Sequence Length etc were calculated. Apart from this, features based on various amino acid descriptor scales like Z3 scale from Hellberg et al. 1987, Eisenberg scale from Eisenberg et al. 1982, GRAVY scale from Kyte and Doolittle 1982 etc were extracted from both positive as well as negative FASTA sequences. Apart from this,amino acid frequency and dipeptide frequency for the FASTA sequences were calculated by using a custom Perl script and stored as a CSV data.

#### Obtaining Top Features

Once the features were extracted from the FASTA sequence datasets, the top features were taken by using scikit-learn (Pedregosa et al., 2011) mutual information gain for selection of top features. Mutual information gain is a measure of the dependency between the features. It is equal to zero if two features are independent of each other, non-zero if there is a dependency between the features. The usage of the mutual information gain is-

```
from sklearn.feature_selection import SelectKBest, mutual_info_classif

selection =  SelectKBest(score_func=mutual_info_classif).fit(X, y)
```

where X is the feature variable and y is a label variable.

For the machine learning model, the features were taken from the amino acid frequency, dipeptide frequency data as well as physicochemical properties like density, molecular weight etc. On conducting validation, it was found that dipeptide frequency and amino acid frequency of the peptides were found to be the best sets of features for the machine learning model. So we extracted 25 best features from a combination of 420 amino acid and dipeptide features and employed them in our final model.

### Building a Machine Learning model

For building the model, We make use of Soft Voting Classification with AdaBoost(Freund and Schapire, 1995) and XGBoost(Chen and Guestrin, 2016) assigned as the estimators for the model, with a high weight assigned to XGBoost classifier model. Adaboost is a meta-estimator algorithm that fits the assigned algorithm to the dataset and any incorrectly classified instances are adjusted for better classification. XGBoost is an implementation of Gradient Boosted trees which have been optimized for speed and accuracy. The model is created using scikit-learn (Pedregosa et al., 2011) and XGBoost(Chen and Guestrin, 2016) Python package.scikit-learnscikit-learn (Pedregosa et al., 2011) and XGBoost(Chen and Guestrin, 2016) Python package.

##### The parameters for the Adaboost model are as follows-

`algorithm="SAMME"` for SAMME discrete boosting algorithm

`learning_rate=0.01` for setting the contribution of the classifier.

`n_estimators=800` for setting the termination number at which boosting will be terminated.

`base_classifier=DecisionTreeClassifier(max_depth=4)` for using Decision Tree as a base classifier with maximum depth as 4.

##### The parameters for the XGBoost model are as follows-

`max_depth=5` for setting the max depth of the Gradient Boosted Tree.

`learning_rate=0.01` for setting the learning rate of the boosted tree.

`min_child_weight=1` for setting minimum sum of hessian weight needed in a child node.

`n_estimators=100` sets the number of Gradient Boosted trees to fit to the dataset.

Both the models were put in a Voting Classifier with a weight of 4:6 with majority in favor of XGBoost Classifier. The accuracy for the model was checked for 5-fold cross-validation and the accuracy comes to around 77.63% and an ROC score of 0.78.

##### Motivation behind selection of the model-

Various models were tested for obtaining the best cross-validation accuracy with the dataset and we found the Ensemble Voting between Adaboost and XGBoost to work really well for the dataset. While we don't

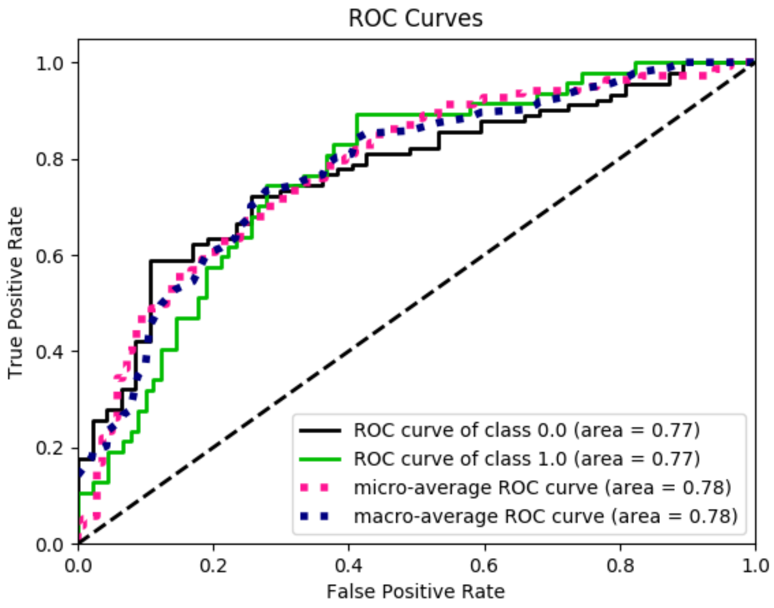


Figure 1: This is a caption

expect improvement in cross-validation accuracy using the ensemble model, we expect to see improvements in accuracy if we make use of the training-testing split.

**Dealing with Imbalance in the dataset-**

To deal with imbalance between positive and negative data points, We have used K-fold cross validation to help keep an equal distribution between the positive and negative classes. We have also made use of Voting classifier of Adaptive and Gradient Boosting, which should help counter the effects of the imbalance in the dataset.

### Biological Relevance of the model

The top features obtained using mutual information gain shows that cysteine-phenylalanine, leucine-leucine, lysine-cysteine were the best dipeptide feature and amino acid frequencies of valine, serine, arginine, asparagine, histidine, cysteine, alanine were the top features which are used for the model and gave a good accuracy for our problem set.

### Code repository for the challenge

The code along with the amino-acid and dipeptide frequency is being hosted at- https://github.com/souravsingh/Ideation-Challenge
