## Supplementary material for "Crowdsourcing Machine Intelligence Solutions to Accelerate Biomedical Science: Lessons learned from a machine intelligence ideation contest to improve the prediction of 3D domain swapping": Model-3_YashShah_Inbix17_paper_61.pdf

INBIX 2017- Ideation Challenge

### Improving the prediction accuracy of 3D domain swapping from sequence information using machine learning


#### Team:

Yash Shah, Thadomal Shahani Engineering College, Bandra-west, Mumbai

Contact no. : +91 9920051150



#### Abstract:

3D domain swapping is a protein structural phenomenon that mediates the formation of the higher order oligomers in a variety of proteins with different structural and functional properties. 3D domain swapping is a mechanism for two or more protein molecules to form a dimer or higher oligomer by exchanging an identical structural element ("domain"). A 5 fold cross validation is performed using feature subsets to improve accuracy learning. The Project classifies protein sequences into 3d Swapped Domain and non 3d swapped using a Neural Network Multi Perceptron Classifier with a Testing Accuracy of 72.5% which can be increased using effective hidden layers and activation functions.


#### Data Summary

Protein Sequences from 3d Swap Knowledgebase Database are used for analysis. The training dataset was constructed using 150 domain swapping and 150 non-domain swapping sequences. Remaining 63 domain swap sequences and 63 non-domain swapping sequences were employed for testing. Neural Network Model is generated using a combination of 66 features derived from sequence, structure and physico-chemical properties.

#### Pre Processing

Dataset consists of 66 features derived from sequence and structural properties of proteins. Due to difference in properties, features have different ranges and scale. In general, learning algorithms benefit from standardization of the data set. If some outliers are present in the set, robust scalers or transformers are more appropriate. Standardization of datasets is a common requirement for many machine learning estimators implemented in scikit-learn; they might behave badly if the individual features do not more or less look like standard normally distributed data. Feature scaling is a method used to standardize the range of independent variables or features of data. In data processing, it is also known as data normalization and is generally performed during the data preprocessing step. The Normalizer rescales the vector for each sample to have unit norm, independently of the distribution of the samples. We have preprocessed our data and scaled in zero mean and unit variation using Standard Scaler library from scikit-learn.

X'= x- mean/ (Standard Deviation).

#### Feature Selection.

Feature Selection is an important Step in Machine Learning to reduce data size and improve accuracy. The feature selection algorithm removes the irrelevant and redundant features from the original dataset to improve the classification accuracy. The feature selections also reduce the dimensionality of the dataset; increase the learning accuracy, improving result comprehensibility. The feature selection avoid over fitting of data. The feature selection also known as attributes selection which is used for best partitioning the data into individual class. Selectkbest method is used for feature selection. **Selectkbest** method is a Univariate feature selection which works by selecting the best features based on univariate statistical tests. It can be seen as a preprocessing step to an estimator. Subset evaluation methods, in contrast, select feature subsets and rank them based on certain evaluation criteria and hence are more efficient in removing redundant features. **SelectKBest** method removes all but the k highest scoring features using a scoring function as statistical test. The methods based on F-test estimate the degree of linear dependency between two random variables.

The Top 10 features selected are

1. Frequency of amino acid "E"
2. 'Frequency of helix'
3. 'Frequency of coil'
4. 'Frequency of solvent inaccessible residues in helix'
5. 'Frequency of solvent inaccessible residues in coil'

6. 'Number of cysteins (invole in disulfide bond formation) in strand'
7. 'hydrogen bond (sidechain to mainchain CO) in helix'
8. 'hydrogen bond (side chain to main chain NH) in helix'
9. Refractivity
10. side chain volume

5 fold cross validation is performed using 5 feature sets, 10, 25,40,55,66.

#### Model Summary:

Multi-layer Perceptron (MLP) is a supervised learning algorithm that learns a function by training on a dataset, where m is the number of dimensions for input and o is the number of dimensions for output. Given a set of features X = {x_1, x_2, ..., x_m} and a target y, it can learn a non-linear function approximated for either classification or regression. It is different from logistic regression, in that between the input and the output layer, there can be one or more non-linear layers, called hidden layers.

Class MLPClassifier implements a multi-layer perceptron (MLP) algorithm that trains using Backpropagation in feed forward networks

MLP trains on two arrays: array X of size (n_samples, n_features), which holds the training samples represented as floating point feature vectors; and array y of size (n_samples,), which holds the target values (class labels) for the training samples.

Neural Network model implements back propagation for error correction on weights on hidden layers. Each neuron in the hidden layer transforms the values from the previous layer with a weighted linear summation w_1x_1 + w_2x_2 + ... + w_mx_m, followed by a non-linear activation function. The output layer receives the values from the last hidden layer and transforms them into output values.

MLPClassifier implements various activation functions and solvers for supervised learning of neural network

**Activation function** for the hidden layer.

‘identity’, no-op activation, useful to implement linear bottleneck, returns f(x) = x

‘logistic’, the logistic sigmoid function, returns f(x) = 1 / (1 + exp(-x)).

‘tanh’, the hyperbolic tan function, returns f(x) = tanh(x).

‘relu’, the rectified linear unit function, returns f(x) = max(0, x)

The **solver** for weight optimization.

'lbfgs' is an optimizer in the family of quasi-Newton methods.

'sgd' refers to stochastic gradient descent.

'adam' refers to a stochastic gradient-based optimizer

Note: The default solver 'adam' works pretty well on relatively large datasets (with thousands of training samples or more) in terms of both training time and validation score. For small datasets, however, 'lbfgs' can converge faster and perform better.

Number of Hidden Layers also matter for accurate results and effective weight computation.

#### Results

Neural Network Model classified the protein sequences into domain swapped and non-domain swapped proteins. The model was trained on a training dataset containing 150 proteins from the positive dataset and 150 proteins from the negative dataset. The performance of the model was evaluated using the five-fold cross-validation method. As shown in Table 2, overall prediction accuracy of 76.67% was obtained by five-fold cross validation.

| Feature Subset | Accuracy Score |
|---|---|
| 10 | 70 |
| 25 | 69.6 |
| 40 | 71.33 |
| 55 | 74.5 |
| 66 | 76.67 |

Model classified data from Test Set with **72.5%** accuracy. This accuracy is obtained using relu activation function and hidden layer of (10,5) size which represents 10 neurons in 1$^{st}$ Hidden layer and 5 neurons in 2$^{nd}$ Hidden Layer.

Accuracy can be increased using various combinations of hidden layers, activation functions and solvers from MLPClassifier.

**Advantages and Biological Relevance of Model**

ANN is nonlinear model that is easy to use and understand compared to statistical methods. ANN is non-parametric model while most of statistical methods are parametric model that need higher background of statistic. ANN with Back propagation (BP) learning algorithm is widely used in solving various classification and forecasting problems. It works for both linear and non linear data. Artificial Neural Network is used for Supervised as well as unsupervised learning. Error Correction is also effective in backpropagation algorithm.

The advantages of Multi-layer Perceptron are:

Capability to learn non-linear models.

Capability to learn models in real-time (on-line learning) using partial_fit.

**LINK TO Github source code**

https://github.com/shahyash-95/ideation.challenge_inbix2017.git
